# *De novo* assembly, delivery and expression of a 101 kb human gene in mouse cells

**DOI:** 10.1101/423426

**Authors:** Leslie A. Mitchell, Laura H. McCulloch, Sudarshan Pinglay, Henri Berger, Nazario Bosco, Ran Brosh, Milica Bulajic, Emily Huang, Megan S. Hogan, James A. Martin, Esteban O. Mazzoni, Teresa Davoli, Matthew T. Maurano, Jef D. Boeke

## Abstract

Design and large-scale synthesis of DNA has been applied to the functional study of viral and microbial genomes. New and expanded technology development is required to unlock the transformative potential of such bottom-up approaches to the study of larger mammalian genomes. Two major challenges include assembling and delivering long DNA sequences. Here we describe a pipeline for *de novo* DNA assembly and delivery that enables functional evaluation of mammalian genes on the length scale of 100 kb. The DNA assembly step is supported by an integrated robotic workcell. We assembled the 101 kb human *HPRT1* gene in yeast, delivered it to mouse embryonic stem cells, and showed expression of the human protein from its full-length gene. This pipeline provides a framework for producing systematic, designer variants of any mammalian gene locus for functional evaluation in cells.

**Significance Statement:** Mammalian genomes consist of a tiny proportion of relatively well-characterized coding regions and vast swaths of poorly characterized “dark matter” containing critical but much less well-defined regulatory sequences. Given the dominant role of noncoding DNA in common human diseases and traits, the interconnectivity of regulatory elements, and the importance of genomic context, *de novo* design, assembly, and delivery can enable large-scale manipulation of these elements on a locus scale. Here we outline a pipeline for *de novo* assembly, delivery and expression of mammalian genes replete with native regulatory sequences. We expect this pipeline will be useful for dissecting the function of non-coding sequence variation in mammalian genomes.

## Introduction

Synthetic genomics is an emerging field focused on building genomes or particularly critical specific segments of them from the ground up. The general impetus is to build designer sequences to specification in order to test and expand our knowledge of biological principles. Beyond the pursuit of knowledge, the ability to write custom genomes with designed characteristics and functions will open doors to solve outstanding problems in healthcare, energy, materials, chemicals, the environment and beyond.

Genome writing projects undertaken to date include viral and microbial genome sequences. In 2002, the 7.5 kb poliovirus was produced by chemical synthesis (1), and since then a number of increasingly large viral genomes in the size range of 100-200 kb have been synthesized (2–4). In 2010, a synthetic version of the bacterial *Mycoplasma mycoides* 1 Mb genome was assembled and shown to be functional following genome transplant into a closely related species (5). Subsequently a *M. mycoides* genome sequence, minimized in size by nearly half through design and complete synthesis, successfully powered growth of a cell (6). The extremely ambitious Synthetic Yeast Genome Project, or Sc2.0, now aims to produce a synthetic genome for the eukaryotic model organism *Saccharomyces cerevisiae.* Distinct from most genome synthesis projects, the goal of Sc2.0 is to write a dramatically modified version of the 12 Mb genome in order to be able to address otherwise unapproachable biological questions. Thus far multiple synthetic chromosomes have been completed (7–14) along with widespread use of the semi-synthetic cells to investigate genome structure-function relationship using the inducible evolution system SCRaMbLE (synthetic chromosome rearrangements and modification by loxP-mediated evolution) (15–20).

The international GP-Write Consortium now aims to define a roadmap and initiate a global effort to responsibly build mammalian and other large genomes from the ground up (21). While substantial technology development will be required to achieve this, ongoing microbial genome writing projects provide many lessons. For instance, a common denominator is the requirement for long DNA sequences that can be readily delivered to the appropriate destination cell. Commercial gene synthesis companies commonly produce 3-5 kb fragments and therefore hierarchical assembly strategies are required to assemble longer sequences. To consider writing mammalian genomes or even segments of them will require massively scaled DNA assembly and the use of automation platforms. In the context of Sc2.0, we have developed strategies for producing synthetic yeast chromosomes (10) that are highly adaptable to assembling any arbitrary DNA sequences, including segments of mammalian genomes. DNA delivery to mammalian cells, on the other hand, requires new technology development to scale and make routine the delivery of DNA at relevant length scales. Current approaches dependent on site-specific integration and landing pads resident in destination cell lines provide an excellent starting point for precise and single-copy integration of designer DNA (22–24).

Here we outline a pipeline for *de novo* assembly, delivery and expression of mammalian genes replete with native regulatory sequences. We describe extrachromosomal Switching Auxotrophies Progressively by Integration (eSwAP-In), for the *de novo* assembly of arbitrary DNA sequence in yeast that allows for purification of large quantities of that material *in vitro.* We demonstrate assembly of a 101 kb human gene, *HPRT1*, and describe an automated system that supports rapid identification of yeast assemblies carrying correctly assembled constructs. Finally, we show that a 114 kb sequence assembled *de novo* in yeast can be delivered and functionally validated in mouse embryonic stem cells. Together this study describes a pipeline that can be applied to the functional evaluation of mammalian gene loci using synthetic genomics. In particular, we anticipate that this pipeline may be useful for dissecting the function of non-coding sequence variation in mammalian genomes.

## Results

### Assembly of long DNA sequences in yeast by eSwAP-In

To assemble long DNA sequences, we devised an assembly strategy called <u>e</u>xtrachromosomal <u>Sw</u>itching <u>A</u>uxotrophies <u>P</u>rogressively by <u>In</u>tegration (eSwAP-In) that uses an *S. cerevisiae* - *E.coli* shuttle BAC vector as a cloning vehicle. The method relies on the inherent capacity of *S. cerevisiae* to perform homologous recombination (HR), which can be harnessed for the process of *‘in yeasto’* DNA assembly. With a minimum of 40 base pairs of terminal sequence homology encoded by adjacent parts, *S. cerevisiae* can stitch together DNA sequences of interest with high fidelity (25–28).

eSwAP-In is a variant of the previously described “SwAP-In” method, which has been used to replace native yeast chromosomal DNA with synthetic designer sequences for the Sc2.0 project (8–10). Both eSwAP-In and SwAP-In rely on iterative use of two different auxotrophic markers embedded near the rightmost ends of DNA segments that are designed for stepwise assembly. Each successive round of *‘in yeasto’* assembly overwrites the previously introduced selectable marker, enabling marker recycling as the length of assembled DNA construct (called an “assemblon”) increases progressively (Figure 1A). The major distinguishing feature of eSwAP-In is that the incoming DNA is assembled extrachromosomally in a circular format and thus replicates and segregates independently of the sixteen native yeast chromosomes. The circular format specified by eSwAP-In is advantageous as the assembled DNA molecule can theoretically be directly recovered into *E. coli*, and large quantities of purified DNA prepared for delivery to the destination organism of choice. To this end, we designed an eSwAP-In “shuttle vector” encoding features to support replication, segregation, and selection in both yeast and *E. coli* (Figure 1B). Importantly, parts that enable delivery to the destination organism of choice can be encoded in the DNA for assembly at the design stage, and thus the eSwAP-In vector may be used universally for assembly of any DNA of interest. To ensure replication and stability of larger eSwAP-In assemblons once transferred into *E. coli*, the eSwAP-In shuttle vector encodes a single copy origin of replication for *E. coli* (bacterial artificial chromosome (BAC)).

**Figure 1:**
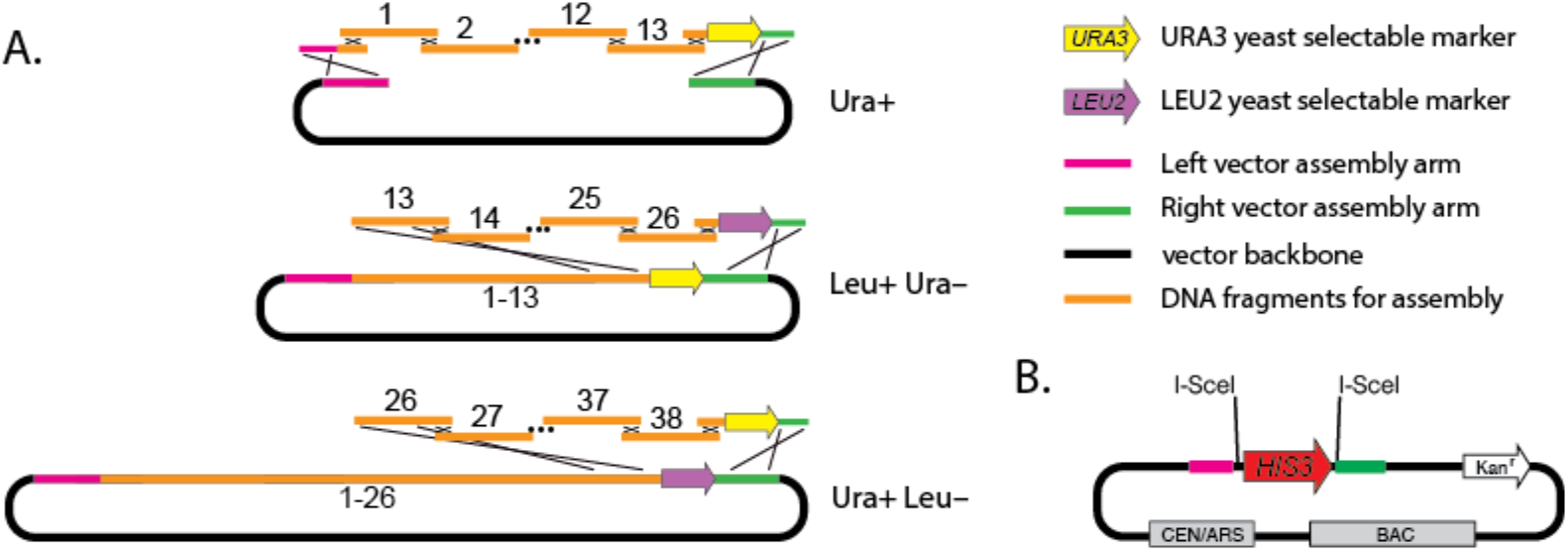
eSwAP-In schematic. **(A)** eSwAP-In *‘in yeasto’* assembly strategy. Three iterative steps of eSwAP-In are shown, switching between yeast selection markers *URA3* (yellow) and *LEU2* (purple) to assemble a circular, extrachromosomal DNA molecule in yeast. Homologous recombination (black X’s) between adjacent parts for assembly (orange), linker sequences, and the vector assembly arms (pink and green) is achieved using a minimum of 80 base pairs of terminal sequence homology. Expected yeast plate-based phenotypes (Leu^+^ Ura^−^, etc.) are indicated. **(B)** The eSwAP-In assembly vector carries parts for segregation and replication in yeast (centromere (CEN), autonomously replicating sequence (ARS)) and *E. coli* (bacterial artificial chromosome (BAC)) so it can be shuttle between the two species. Kanamycin (Kan^r^) is used for selection in *E. coli.* The vector can be linearized *in vitro* or in yeast using I-*Sce*I to drop out the yeast selection marker *HIS3.*

### *HPRT1* assembly by eSwAP-In

The use of eSwAP-In to assemble long DNA sequences can be readily applied to the study of mammalian gene loci. Mammalian genomes consist of a tiny proportion of relatively well-characterized coding regions and vast swaths of poorly characterized “dark matter” containing critical but much less well defined regulatory sequences; *de novo* assembly of mammalian gene loci and designer versions of those loci using eSwAP-In can be used to dissect the function of non-coding regions. *De novo* assembly of a given gene locus is also advantageous as delivery can be made to ectopic loci in the target genome or even to cells of different species, enabling gene functional evaluation that goes well beyond the limits of CRISPR-Cas9 editing.

To demonstrate eSwAP-In, we set out to build a ~100 kb human gene locus, and chose the highly conserved human hypoxanthine-guanine phosphoribosyltransferase (*HPRT1*) gene (Fig. 2A). *HPRT1* encodes an enzyme with a key role in purine salvage and is well studied as it is used in cell culture both as a selectable and counter-selectable marker. Moreover, mutations in this well studied gene, encoding a critical enzyme in the purine salvage pathway, underlie a series of human inherited conditions ranging from gout to Lesch-Nyhan syndrome (29, 30). Using three sequential steps of eSwAP-In, we assembled the ~101 kb wild type human *HPRT1* (*hHPRT*) locus in yeast. This was achieved by producing 38 x ~3kb PCR amplicons spanning the length of *hHPRT1* (Fig. 2B) using human genomic DNA extracted from HEK293T cells as template. Each amplicon encodes >80 bp of homology at each end with adjacent assembly parts (Fig. 2B), providing substrates for homologous recombination between adjacent parts for assembly in yeast. In three sequential eSwAP-In assembly steps, we assembled ~35-40 kb of the gene locus using 12 or 13 PCR amplicons at each stage (Fig. 2C, 2D) plus the appropriate terminal “linker” segments (see Materials and Methods). Each successive round of *‘in yeasto’* assembly overwrote the previously introduced selection marker (auxotrophy), enabling selection marker recycling as the assemblon grew (Fig. 2D). For each assembly step, we screened yeast transformants with the correct marker phenotypes (e.g. Ura^+^/Leu^−^) for the presence of assembly junctions using spanning primers (described in detail below). Assemblons and intermediates built by eSwAP-In were easily recovered from yeast into *E. coli* for digestion verification (Fig. 2E). We efficiently assembled a 101 kb human gene locus based on the wild-type *hHPRT1* in three eSwAP-In steps in about six weeks.

**Figure 2:**
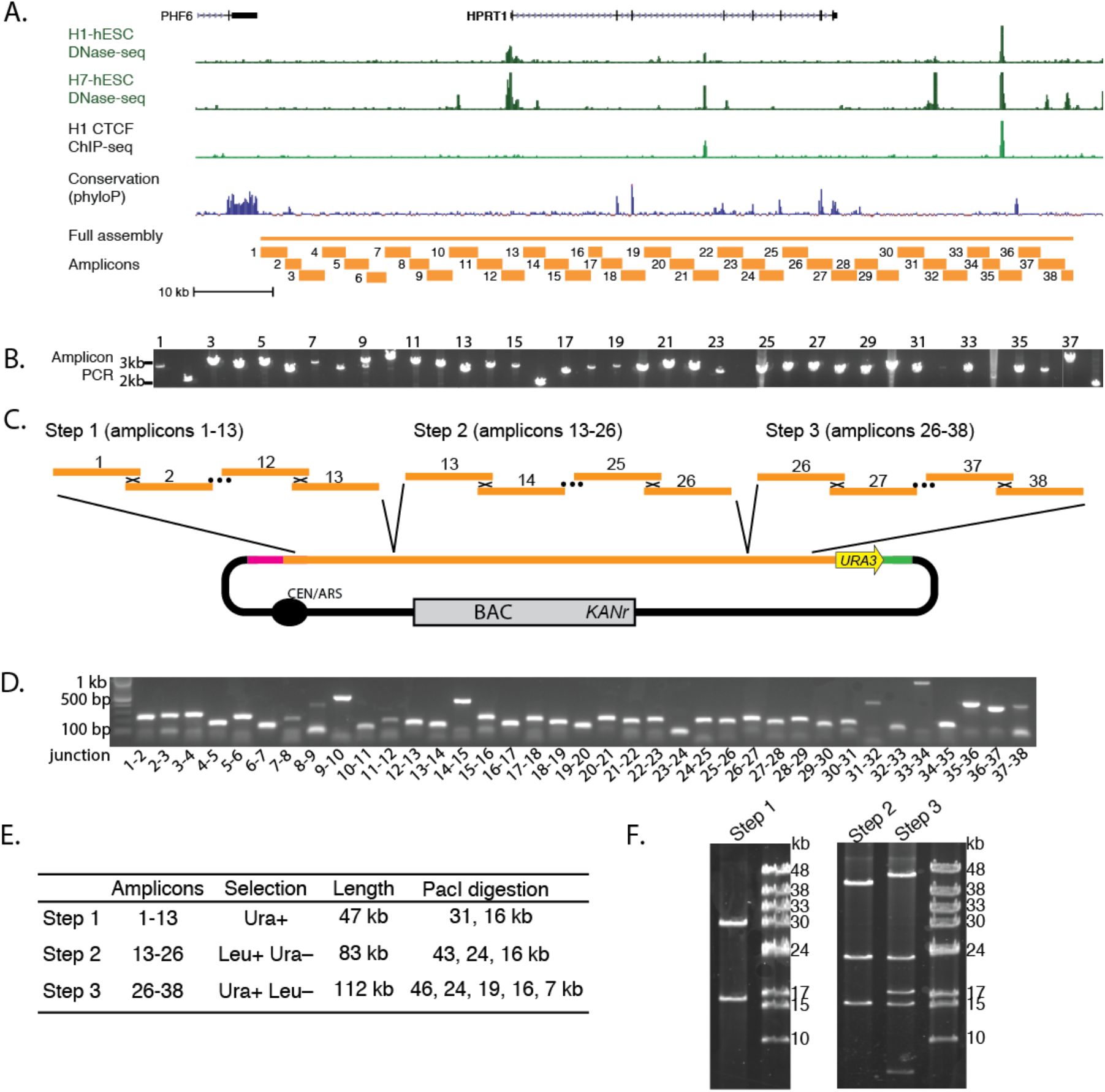
Assembly of 101 kb *hHPRT1* gene locus by eSwAP-In. **(A)** *hHPRT1* locus showing CTCF binding sites and DNase hypersensitive elements along with conservation. Amplicons (~3 kb; orange lines) were designed to tile the locus and overlap each other by at least 80 bp for assembly in yeast. **(B)** Verification gel showing 38 tiled amplicons used for eSwAP-In assembly. **(C-D)** 3-step eSwAP-In assembly strategy. The construct is flanked by Not*I* sites at either end. Length includes the 11 kb vector. **(E)** Intermediate (step 1 (pLM718) and 2 (pLM747)) and final (step 3 (pLM750)) assemblons were recovered into *E. coli* for *Pac*I restriction digestion verification using field inversion gel electrophoresis (FIGE).

The seαuence of the full-lenεth *hHPRT1* construct was determined using Pacific BioSystems long read sequencing. Relative to the human reference genome sequence, we uncovered 66 sequence variants, including 41 substitutions and 25 deletions. This tally excludes length variations associated with homopolymer runs (five or more identical bases neighboring one another), which could originate from misannotations in the reference sequence, errors in PCR, variations during yeast assembly, or artifacts of sequence analysis. The 66 variants are likely to represent a combination of PCR-induced errors and bona fide differences from the reference in the HEK293T genome, which was used as the template DNA for producing the amplicons. None of the variants were in *HPRT1* coding exons.

### Efficiency of assembly - automated screening of yeast transformants

A critical parameter affecting eSwAP-In and the time it takes to assemble a given gene is the number of amplicons that can be simultaneously assembled in one step in yeast. For example, increased multiplexing of assembly fragments will reduce the number of intermediate eSwAP-In steps, but requires additional screening to identify correctly assembled transformants. We devised a simple genotyping assay to evaluate the structure of DNA assembled in yeast transformants. Specifically, we evaluate the presence or absence of PCR amplicons produced from primer pairs spanning regions of terminal homology, a.k.a. “junctions”, between adjacent assembly fragments (Fig. 3A). The presence of all junction amplicons is consistent with correctly assembled constructs in yeast.

**Figure 3:**
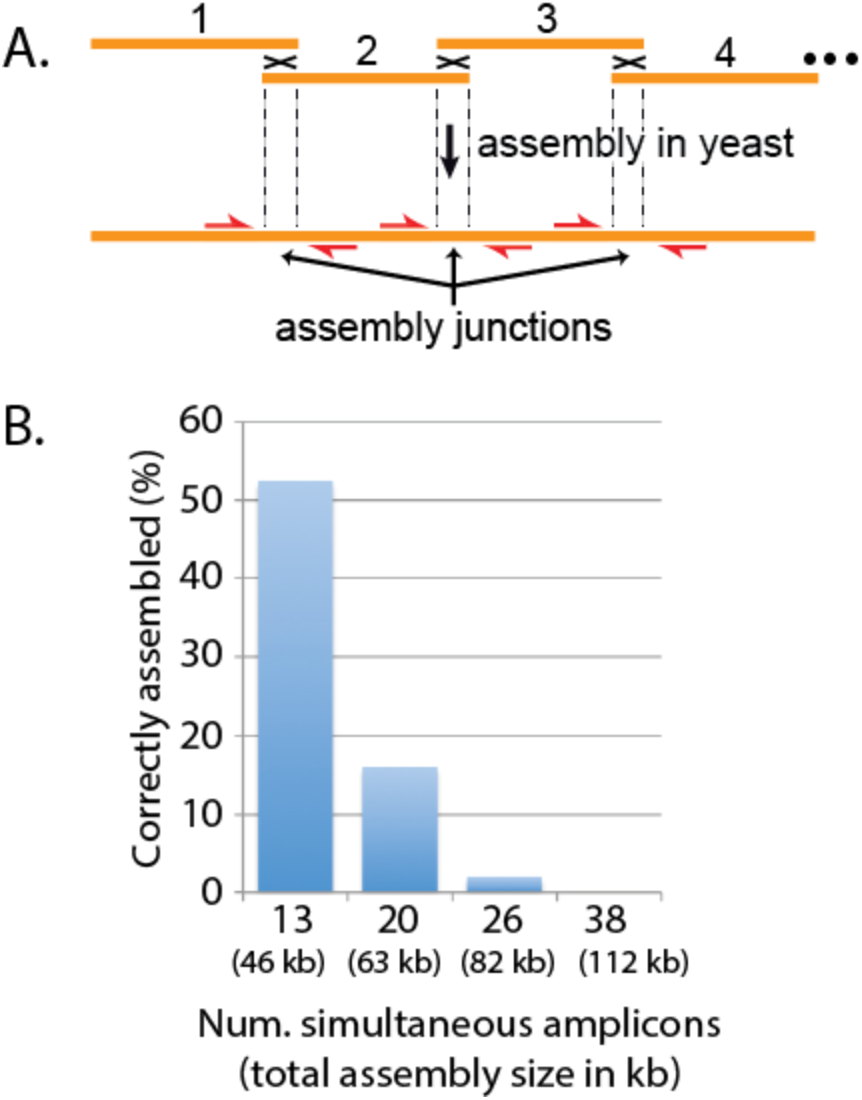
Automated high throughput junction PCR analysis of 511 single-step assemblies. **(A)** Primers (red arrows) spanning assembly junctions test for the presence/absence of amplicons in many independent yeast colonies. Reactions are performed using the automated workcell in a 500 nL volume. **(B)** Assembly success (the fraction of total colonies tested with all assembly junctions intact) as a function of the number of PCR amplicons simultaneously transformed into yeast for assembly.

We previously described a high throughput genotyping workflow for the Sc2.0 synthetic yeast genome project (31). To enable high throughput screening of assembly junctions, we integrated this workflow into a robotic workcell capable of testing >30,000 assembly junctions per day. The workcell includes robotic arms to transfer source and destination plates between plate hotels and nests, a bulk liquid dispenser, an acoustic droplet ejection (ADE) robot, a centrifuge, a plate sealer, and a 1536 well real time PCR machine. In brief, after dispensing 500 nL of PCR mastermix to each well of a 1536 plate, 10 nL each of primers and template DNA are distributed to specified wells by the ADE robot, and the plate is spun down, sealed and inserted into the 1536 thermal cycler. Template DNA derives from independent yeast transformants of a given assembly, which are first re-arrayed for liquid growth in selectable medium in a 96 deep-well plate by a colony picker, and later processed for template DNA preparation by modified alkaline lysis and neutralization. Primer pairs that span assembly junctions are tested in individual PCR reactions against each template DNA preparation. ‘Winners’ are defined as colonies that produce amplicons across all assembly junctions.

To begin to understand the relationship between the number of amplicons and assembly efficiency, we used the workcell to characterize assembly junctions of 511 assemblons derived from different numbers of assembly fragments (Fig 3). We used reagents from the *HPRT1* project, attempting assembly of 13, 20, 26, or 38 PCR amplicons in a single yeast transformation. In all cases, the same left linker was used to provide homology between the first PCR amplicon and the left end of the linear assembly vector. An assembly-specific right linker was either re-used from the *HPRT1* eSwAP-In assembly (13, 26, and 38 fragment assemblies) or generated for this purpose (20 fragment assembly) to provide homology between the rightmost PCR amplicons and the other end of the vector. Our results indicate our ability to assemble 47 kb constructs from 13 PCR amplicons plus two linkers and an assembly vector, effectively a 16 piece assembly, with >50% success (Fig 3B). Assembly efficiency declines with increasing numbers of PCR amplicons, dropping to 16% for 20 fragments and 3% for 26 fragments; despite screening >100 yeast colonies, we did not find any transformants with all assembly junctions intact for 38 fragments.

### Delivery of *hHPRT1* to mouse embryonic stem cells

To enable delivery of large DNA constructs to mammalian cells we made use of a previously described system, Inducible Cassette Exchange (ICE), which was previously utilized to deliver ~10 kb constructs to mouse embryonic stem cells (mESCs) with high efficiency (23, 32). The ICE-enabled mESC line (A17iCre) carries a “landing pad” on the X chromosome that encodes a doxycycline-inducible *CRE* transgene flanked by heterotypic, self-incompatible lox sites (Fig. 4A). Downstream of the floxed *CRE* is a G418 resistance gene lacking a start codon and promoter. Treating cells with doxycycline causes Cre to be expressed (rtTA is inserted into the Rosa26 locus), rendering cells recombination-competent. Delivery of DNA carrying the same heterotypic lox sites results in cassette exchange recombination and the replacement of CRE with the incoming DNA. Recombinants are selected based on neomycin resistance (G418^R^) and GFP expression in the presence of doxycycline.

**Figure 4:**
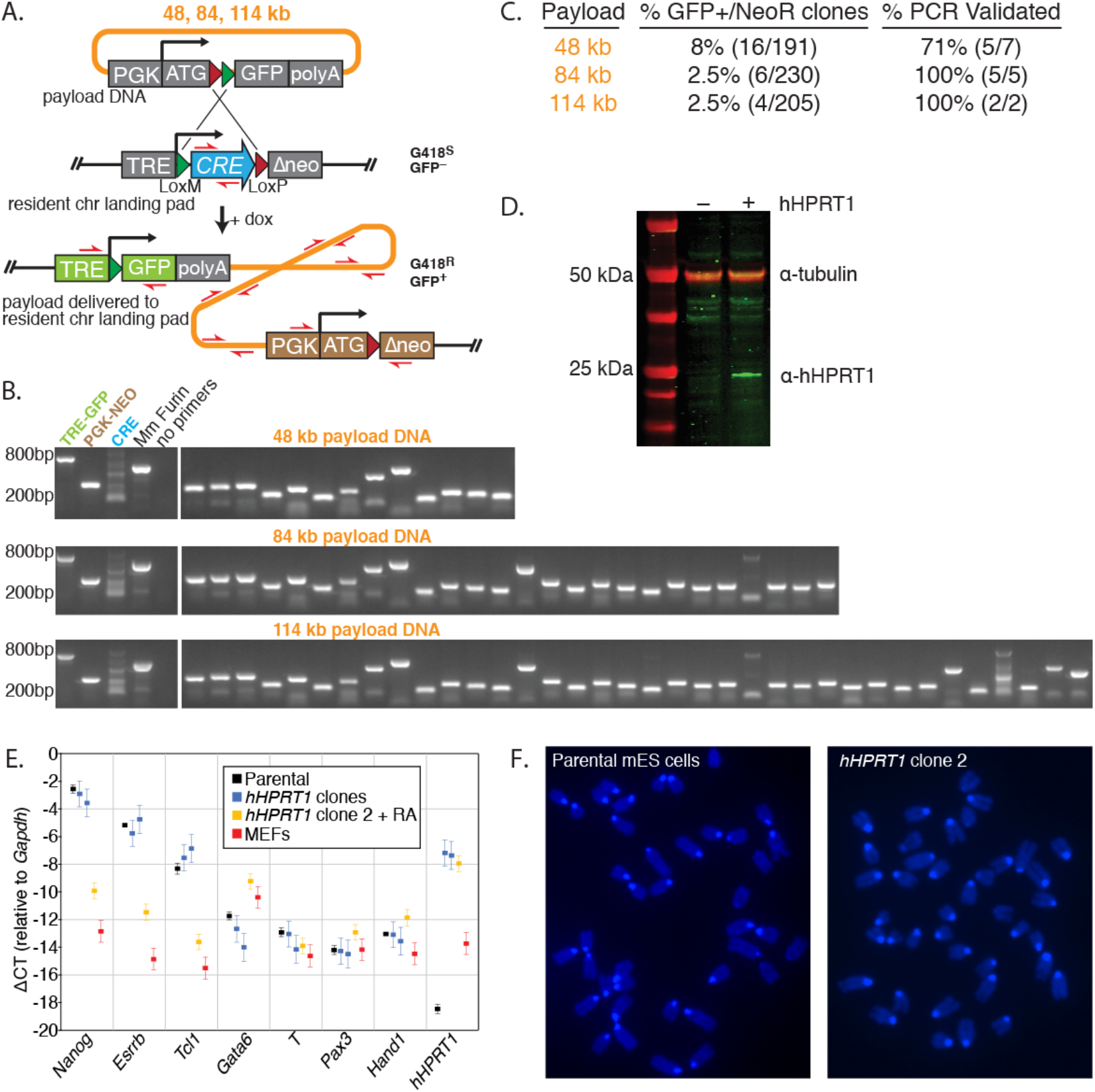
Delivery of big DNA by ICE and clone characterization. **(A)** Schematic of integration of DNA into a resident “landing pad” in a mouse ES cell line using ICE. Integrants are selected for G418 resistance and GFP expression in the presence of doxycycline (+ dox). In this system, the rtTA gene is separately expressed from the *Rosa26* locus. In this experiment, incoming DNA for delivery included the *HPRT1* partial or full-length assemblons (37, 71, or 101 kb), the loxP-loxM cassette (~2 kb), and the vector backbone (~11 kb). **(B)** PCR analysis of independent clones from the delivery of 48 kb, 84 kb and 114 kb constructs. Primer pairs (red) span the length of the delivered DNA, as well as newly-formed junctions post-integration. **(C)** Correct delivery frequency reported as a fraction of clones that are both GFP positive and neomycin resistant as compared to the total number of neomycin resistant clones. PCR validation as in (B). **(D)** Western blot analysis for *hHPRT1* protein expression in mouse ES cells using a human HPRT-specific monoclonal antibody. Parental mES (−) and *hHPRT1* clone 1 (+) were used. Similar results were obtained for *hHPRT1* clone 2 (See Supplemental Fig. 1). **(E)** Quantitative real-time PCR analysis of the parental mESCs and the two clones carrying *hHPRT1.* Data are presented as mean ΔCT +/− S.D. relative to *Gapdh.* Control samples include retinoic acid-treated mES *hHPRT1* clone 2 (+RA) and immortalized mouse embryonic fibroblasts (MEFs). Measured mRNAs include pluripotency markers *Nanog, Esrrb* and *Tcl1*, endoderm differentiation marker *Gata6*, mesoderm differentiation marker *T* (Brachyury), ectoderm differentiation marker *Pax3*, trophectoderm differentiation marker *Hand1* and *hHPRT1.* **(F)** Representative metaphase spreads from parental mESCs and *hHPRT1* clone 2. Similar results were obtained for *hHPRT1* clone 1 (See Supplemental Fig. 1).

Starting with constructs derived from each of the three *HPRT1* eSwAP-In assembly steps (Fig. 2D), an ICE cassette (PGK-ATG-loxP-loxM-GFP-polyA, ~2 kb) was added at the 5’ end of each assembly (~30 kb upstream of the *HPRT1* start codon), to make them compatible with delivery to the resident chromosome landing pad in the mESCs (Fig. 4A). DNA was purified by cesium chloride/ethidium bromide banding and delivered by nucleofection to mESCs pre-treated with doxycycline. In this experiment, a supercoiled circular molecule consisting of both the vector and insert DNA was delivered to the mESC ICE cassette resulting in total DNA lengths integrated of 48, 84, or 114 kb. GFP-positive and neomycin-resistant clones were evaluated by PCR for the presence of the delivered DNA plus newly-formed junctions between landing pad and genome (TRE-GFP, PGK-NEO) (Figure 4B). The frequency of GFP-positive clone formation and the PCR validation is shown in Figure 4C; almost all GFP positive clones appear to carry the entire payload DNA, suggesting that ICE can be used to faithfully deliver large constructs to mESCs. Overall, we demonstrated successful delivery of 48, 84, and 114 kb DNA constructs to mouse ESCs using ICE.

The two clones derived from delivery of the 114 kb constructs encode the 101 kb full length *hHPRT1* gene (40 kb gene body plus 30 kb flanking regions on each side, plus the vector sequence and ICE cassette). We evaluated protein expression by immunoblot using a monoclonal antibody directed against *hHPRT1.* Expression of the human protein in mouse cells was readily detected in each of two clones tested (Fig. 4D and Supplemental Fig. 1).

To test whether the delivery of large DNA constructs and the selection strategy we employed affected the pluripotency status of our mESCs, we measured the mRNA levels of a series of pluripotency and early differentiation markers (33) using quantitative real-time PCR. As depicted in Figure 4E, the parental mESCs, as well as the two clones carrying the full-length *hHPRT1* assemblon DNA had nearly identical patterns of marker gene expression. Specifically, we observed high levels of the pluripotency markers *Nanog, Esrrb* and *Tcl1* and low expression levels of markers of endoderm (*Gata6*), mesoderm (T), ectoderm (*Pax3*) and trophectoderm (*Hand1*) differentiation. Moreover, robust transcriptional activation of *hHPRT1* was detected in the two *hHPRT1* clones, compared to their parental A17iCre cells, further validating the correct integration of a fully functional and transcriptionally active *hHPRT1* gene. In addition, to exclude the onset of chromosomal instability or aneuploidy during ICE delivery and selection, we analyzed metaphase spreads from the parental mESCs and the two isolated full-length *hHPRT1* clones. DAPI-stained metaphase spreads showed no difference in chromosome number (40,XY) and no evidence of marker chromosomes or gross chromosomal rearrangements in the parental mESCs or the *hHPRT1* clones (Fig.4F and Supplemental Fig. 1). Together these data suggest delivery of large DNA to mESCs by ICE is precise and does not adversely affect pluripotency or genome stability. In summary, we have assembled, delivered, and shown expression of a human protein from its intact full-length gene in mESCs.

## Discussion

Design and synthesis of long DNA sequences is a technology that could spark a revolution in the functional analysis of mammalian genomes. Given the dominant role of noncoding DNA in common human diseases and traits, the interconnectivity of regulatory elements, and the importance of genomic context, *de novo* design, assembly, and delivery can enable large-scale manipulation of these elements on a locus scale.

Here we describe an iterative *‘in yeasto’* DNA assembly strategy, eSwAP-In, and demonstrate its utility by assembling a 101 kb human gene locus *hHPRT1.* Importantly, using the ICE delivery system (23, 32) we show delivery of a 114 kb construct carrying the *hHPRT1* locus to a mouse cell. With this workflow in place, we are now positioned to build a library of sequence variants of the *hHPRT1* locus and begin to dissect the function of regulatory elements, single nucleotide polymorphisms, and parts that specify 3D genomic architecture. Such “synthetic haplotypes” can be assembled in parallel from a combination of PCR amplicons and synthetic DNA fragments.

Despite the recent emergence of CRISPR-Cas9 to enable surgical alteration of individual sites of interest, there still exists a critical gap in our ability to manipulate multiple sequence features on a consistent haplotype in mammalian genomes, as well as to explore the effect of larger rearrangements of regulatory element positioning and context. The *de novo* design, synthesis, and delivery pipeline described here provides an approach to tackling such limitations.

In theory, the pipeline demonstrated for *hHPRT1* can be applied to any gene locus (mammalian or otherwise) of any length. For instance, eSwAP-In can be used to progressively assemble larger DNA constructs in yeast. However, this depends on (i) the ability to source the required DNA for assembly, either by PCR or commercial synthesis; (ii) how well yeast tolerates the sequence composition across the growing assemblon; and (iii) the upper length limit for chromosome stability in yeast. Factors like high GC content and direct repeats can be problematic for PCR, commercial gene synthesis and yeast assembly. Further, yeast origins of replication may need to be encoded in long and/or GC rich assemblies to promote stability in yeast (34), although this would effectively leave unwanted “scars” in the designed sequence. Independent of sequence composition, we expect the upper length limit of bacterial or mammalian constructs assembled and maintained in yeast to be well over 1 Mb, as has been previously reported (5, 35). Our theory is based on recent work demonstrating that yeast tolerates linear chromosomes ranging in length from 6-12 Mb (36, 37), suggesting that DNA in this size range is not too long for accurate replication and segregation at cell division. Increasing the size of assemblons in yeast will require new approaches to efficiently deliver the constructs to mammalian cells, as large constructs will be subject to shear forces when manipulated *in vitro.* Potential strategies for larger DNA delivery include embedding DNA in agarose plugs (38) or employing direct cell fusion approaches (35, 39).

Ultimately, the pipeline described here lays the groundwork for routine design, construction and delivery of increasingly large, megabase-sized DNA sequences to mammalian cells. For widespread adoption beyond our lab, we project that a number of technical advances may facilitate this vision, including continued decreases in commercial gene synthesis cost (supplanting use of PCR), expanded interest in chromosome and genome-scale cell engineering to address systems-level biological questions, and increased availability and adoption of laboratory automation.

## Materials and Methods

### Strains and media

Yeast assembly by eSwAP-In was performed in BY4741 (40) using standard yeast media. ElectroMAX DH10B cells (Invitrogen, 18290015) were used for recovering assembled DNA from yeast into *E. coli.* Recovery of assembled DNA from yeast to *E. coli* was performed as previously described (8) and *E. coli* electroporation carried out as per the manufacturer’s protocol. Small-scale *E. coli* cultures (5-10 mL) for alkaline lysis and digestion verification were grown in LB medium supplemented with 50 μg/ml kanamycin. Large-scale *E. coli* cultures (400-500 ml) for prepping DNA by cesium chloride to be delivered to mammalian cells were grown in LB medium supplemented with 25 μg/mL kanamycin.

### 38 x 3kb *hHPRT1* amplicons

The *hHPRT1* locus was arbitrarily defined as the ~40 kb gene body of *hHPRT1* plus the 30 kb upstream and downstream. The final coordinates selected encompass 134428938-134534487 on the X chromosome (human genome assembly 38). Primers were designed to produce ~3 kb amplicons with terminal overlaps ~180 bp in length on average, but ranging from 80 bp to 830 bp. Primer positions were selected to anneal outside of repetitive sequences across the human genome as identified by RepeatMasker (41). Primer sequences are listed in Supplementary Table 1. Human genomic DNA was isolated from HEK293T cells and used as template in 20 uL PCR reactions using KAPA HiFi HotStart ReadyMix PCR kit with 0.5 μM of each primer and ~100 ng of genomic DNA. Thermal cycling was carried out as per the manufacturer’s instructions with an extension time of 4 minutes. All but a single primer pair produced an amplicon of the expected length. For amplicon 24, an additional set of primers was designed and all four combinations tested (by pairing the new primers with the original pair) using different polymerases and gradient annealing temperatures to identify the conditions for successful amplification.

### Linker fragments

Linkers to provide homology between terminal assembly fragments and the vector backbone were produced by two- or three-piece fusion PCR using Phusion polymerase as per the manufacturer’s instructions (NEB, M0530). The left linker fragment for the first assembly step encoded 250 bp of the vector and 250 bp of the left-most assembly fragment. For subsequent assembly steps the rightmost *HPRT1* PCR amplicon from the previous step was used, effectively over-writing the resident marker from the previous assembly step. The right linker fragments encoded 250 bp of the rightmost assembly fragment (13, 26, or 38), a yeast selectable marker (*URA3* or *LEU2*) and 250 bp of the vector. The *hHPRT1* gene was built from left to right in three sequential steps.

### Three step eSwAP-IN to assemble *hHPRT1* in yeast

Yeast transformations were carried out as previously described (9). For assembly step 1, I-*Sce*I-linearized pLM453 (~100ng) was co-transformed into yeast along with ~100-200 ng of *HPRT1* amplicons 1 through 13, as well as the appropriate left and right linker fragments. Following selection on synthetic complete medium lacking uracil (SC–Ura), 48 independent transformants were subjected to junction PCR analysis using the integrated workcell; 28/48 colonies passed this screen with at least one primer pair producing amplicons for all tested junctions. Two yeast transformants, frozen as yLM1227 and yLM1228, were used directly for assembly step 2 in which *hHPRT1* amplicons 13-26 were co-transformed along with the right linker fragment encoding the *LEU2* selectable marker. Following selection on synthetic complete medium lacking leucine (SC–Leu), transformants were replica plated onto SC–Ura medium to identify Leu^+^/Ura^−^ colonies, 6 of which were subjected to junction PCR analysis. Two out of six colonies tested produced amplicons for junctions 13-26, yLM1229 (derived from yLM1227), and yLM1231 (derived from yLM1228). The final assembly step was carried out using yLM1229 and yLM1231, co-transforming *hHPRT1* amplicons 26-38 together with a right linker fragment encoding the *URA3* gene. Nine Ura+/Leu− colonies were tested by junction PCR analysis and two yeast colonies, yLM1234 and yLM1235, derived from yLM1229 and yLM1231 respectively, were selected for further analysis.

### Junction PCR analysis

Two sets of primer pairs spanning each assembly junction were used to evaluate the structure of assemblons in yeast colonies when using the automated workcell. Briefly, using a QPix (Molecular Devices), yeast colonies were inoculated into 1 mL of the appropriate selective medium (synthetic complete medium lacking uracil or leucine) and grown with orbital shaking at 30°C for 48 hours. As a control, a yeast colony carrying an empty vector was grown in parallel and a single well was left empty to monitor for contamination and to serve as a no DNA control for downstream real-time PCR. Yeast genomic DNA was prepared by alkaline lysis by mixing 20 μL of saturated yeast culture with 70 μL of 0.02 M NaOH, heating at 70°C for 10 min, and neutralization with 60 μL 0.4 M Tris pH 8. All manipulations were carried out in 96 well plate format using a CyBio Felix (AnalytikJena) and genomic DNA was transferred to an Echo Qualified 384-Well Polypropylene Microplate (LabCyte, Inc., PP-0200). Junction primers were ordered from Integrated DNA Technology, pre-mixed at 50 μM each, and arrayed on an Echo Qualified 384-Well Polypropylene Microplate (LabCyte, PP-0200). A Cobra bulk liquid dispenser (Art Robbins Instruments), was used to dispense 500 nL of Lightcycler 1536 DNA Green qPCR master mix (Roche, 05573092001) into each well of a Lightcycler 1536 multiwell plate (Roche, 05358639001). An Echo 550 acoustic droplet ejection robot (LabCyte) was used to dispense 10nL each of primers and yeast genomic DNA into specified wells of the 1536 Lightcycler multiwell plate. All primers pairs were tested against all experimental genomic DNA samples and controls. A single control primer pair targeting the eSwAP-In assembly vector backbone was used as a positive control to evaluate the quality of each genomic DNA template. Thermal cycling was carried out in the Lightcycler 1536 (Roche). Lightcycler data was analyzed using the MaxRatio method (42). Yeast colonies were considered ‘winners’ if a minimum of one junction primer pair produced an amplicon.

### Structural and sequence verification of assembled DNA

Junction PCR-verified assembled constructs from yLM1227, yLM1228, yLM1229, yLM1231, yLM1234, and yLM1235 (Supplementary Table 2) were all recovered into *E. coli* as previously described (8). The resulting plasmids, pLM718, pLM719, pLM747, pLM749, pLM750, and pLM751 (Supplementary Table 3), respectively, were prepped by alkaline lysis and isopropanol precipitation from 5 mL of saturated overnight culture and subjected to digestion using *Pac*I. Digestion products were separated by field inversion gel electrophoresis (FIGE) using the auto-algorithm of the CHEF Gel mapper (Biorad) and the Monocut ladder (NEB). pLM750 was subjected to Pacbio sequence verification.

### Retrofitting constructs with ICE cassette

*hHPRT1* eSwAP-In Step 1 (pLM718), Step 2 (pLM747) and Step 3 (pLM750) constructs were digested with *Not*I, which either linearized (pLM718, pLM747) or dropped out the insert (pLM750). The ICE cassette was amplified from a pre-existing plasmid (pLM707) using overhang primers that produced 40 bp terminal homology on either side of the leftmost *Not*I site in the three constructs. The ICE cassette PCR product along with linear pLM718 and pLM747 were co-transformed into yeast for assembly. The ICE cassette PCR product along with the right linker and the double digested pLM750 were co-transformed into yeast for assembly. Assemblies were recovered into *E. coli* and sizes verified by digestion and FIGE. Sanger sequencing was performed across the *PGK1* promoter of the ICE cassette and a sequence perfect clone was identified in all cases to produce pLM854, pLM848, pLM881 and pLM886 (Supplementary Table 3).

### mESC culturing conditions

mESCs were grown in ‘80/20’ medium, a mix of 80% 2i medium and 20% mESC medium. 2i medium was made from a 1:1 mix of Advanced DMEM/F12 (Gibco 12634010) and Neurobasal-A (Gibco 10888022), containing 1X N-2 supplement (Gibco 17502048), 1X B-27 supplement (Gibco 17504044), 2mM Glutamax (Gibco 35050061), 0.1mM Beta-Mercaptoethanol (Gibco 31350010), 10^3^ units/mL LIF (Millipore, ESG1107), 1 μM MEK1/2 inhibitor (Stemgent, PD0325901), and 3 μM GSK3 inhibitor (R&D Systems, CHIR99021). mESC medium was made from Knockout DMEM (Gibco 10829018), containing 15% Fetal Bovine Serum (Gemini), 0.1 mM Beta-Mercaptoethanol (Gibco 31350010), 1X MEM Non Essential Amino Acids (Gibco 11140050), Glutamax (Gibco 35050061), 1X Nucleosides (Millipore ES-008-D) and LIF (Millipore ESG1107).

### DNA delivery

DNA (pLM848, pLM881, pLM886) was prepared for delivery to mESCs by cesium chloride (CsCl) banding as described in the Molecular Cloning Laboratory Manuals following Protocol 3 (Preparation of Plasmid DNA by Alkaline Lysis with SDS: Maxipreparation), Protocol 10 (Purification of Closed Circular DNA by Equilibrium Centrifugation in CsCl-Ethidium Bromide Gradients: Continuous Gradients), and Protocol 12 (Removal of Ethidium Bromide from DNA by Extraction with Organic Solvents) (9). Briefly, a 5 mL starter culture, grown in LB kanamycin (25 μg/ml) at 30°C for 24 hours, was used to inoculate 400-500mL of LB kanamycin (25 μg/mL) cultures, which were grown for 24 hours at 30°C. Cell pellets were always frozen at −20°C prior to alkaline lysis and DNA precipitation. CsCl banding was performed in a L-80 ultracentrifuge (Beckman) at 20°C for 16 hours at 45,000 rpm in a VTi 65 rotor (Beckman). Ethidium bromide was extracted from the CsCl banded DNA using water-saturated n-butanol. Purified DNA was resuspended in 12 μL TE + 50 mM NaCl or water. 1μL of CsCl DNA preps was used for digestion and FIGE, and the remainder was used for mESC transfection. The concentration of DNA was determined from the FIGE band intensity relative to the known quantity in the ladder (NEB, N3019S) and approximately 10-20 μg of CsCl prepped DNA was used for each delivery experiment.

DNA was delivered to A17iCre mESCs (23, 32) by nucleofection (Amaxa Nucleofector 2b, mESC Nucleofector Kit, Lonza, VVPH-1001). mESCs were treated with 1 μg/mL doxycycline for 18 hours to induce Cre recombinase. Cells were then harvested using Accutase (Biolegene, 423201) and 5 million cells were nucleofected using program A-023 as per the manufacturer’s instructions. Cells were plated onto gelatinized (0.1% gelatin) 10 cm plates. Selection with 400 μg/mL Geneticin (Life Technologies, 10131-027) was applied 24 h after nucleofection. GFP-positive clones were identified following induction with 3 μg/mL doxycycline and picked 7-10 d post selection. Genomic DNA was extracted from picked clones using the DNeasy Blood & Tissue Kit (Qiagen, 69504) and screened using PCR using primers spanning the assemblon length, as well as newly-formed junctions with the genome. Independent delivery experiments using DNA prepped on separate days was performed for all three constructs. The frequency of GFP positive clones shown in Fig. 4C is from a single experiment but representative of multiple experiments.

### Testing *hHPRT1* expression in mouse cells by immunoblot

Approximately 5 million cells from each positive pLM886-delivered mESC clones and parental A17iCre mESCs were lysed in SKL Triton lysis buffer (50 mM HEPES pH7.5, 150 mM NaCl, 1 mM EDTA, 1 mM EGTA, 10% glycerol, 1% Triton X-100, 25 mM NaF, 10 μM ZnCl_2_) supplemented with protease inhibitors (Sigma, 11873580001). Equal amounts of protein (as measured by a Bradford assay) were loaded onto 4-12% BisTris polyacrylamide gels for separation by electrophoresis. Proteins were transferred onto an Immobilon-FL membrane (Millipore), blocked for 1 hr with blocking buffer (LiCOR) and then probed with rabbit anti-Tubulin and mouse anti-hHPRT antibodies (mybiosource.com, MBS200197). Bands were visualized with LiCOR fluorescent Goat anti-mouse and anti-rabbit secondary antibodies using an Odyssey CLx scanner (LiCOR).

### Quantitative real-time PCR (QRT-PCR)

Total RNA was extracted using the RNeasy kit (QIAGEN). 2 μg RNA were reverse transcribed using the SuperScript IV Reverse Transcriptase kit (ThermoFisher Scientific, 18091200)) according to the manufacturer’s protocol. QRT–PCR was performed in quadruplicates using the KAPA SYBR FAST qPCR Master Mix (KAPA BIOSYSTEMS, KK4610) on a LightCycler480 Real-Time PCR System (Roche) according to manufacturers’ protocols. Expression was calculated using the ΔCT method relative to the expression of the housekeeping gene *Gapdh.* A CT value of 40 was assigned to undetectable samples. Gene-specific primers are listed in Supplementary Table 4. As negative controls for pluripotent mESC expression profiles we analyzed retinoic acid (RA; Tocris Biosciences, 0695)–treated mESCs and immortalized mouse embryonic fibroblasts (MEFs). RA treatment was performed for one of the *hHPRT1* clones by growing the cells in mESC medium lacking LIF and supplemented with 5 μM RA for two days.

### mESC metaphase analysis

For chromosome preparation, cells at ~70% confluency were treated with 0.4 μg/mL colcemid (Roche, 10295892001) for 2 hours. Cells were harvested by trypsinization and both supernatant and PBS wash collected by centrifugation (300 x g, 2 minutes). Pellets were washed in PBS and swollen with 10 ml of 75 mM KCl, pre-warmed to 37°C. Following incubation at room temperature for 5 min, cells were pre-fixed with 500 μL of fixative solution (3:1 methanol:acetic acid, ice cold) and incubated at room temperature for an additional five minutes. Cell pellets were collected by centrifugation (300 x g, 2 min) and the supernatant decanted, saving ~500 μL in which cells were resuspended. 10 mL of fixative solution was slowly added to each sample while gently vortexing. Cell pellets were immediately collected by centrifugation (300 x g, 2 min), supernatant decanted and cells resuspended in ~200-500 μL of residual fixative solution. Samples were stored on ice or at 4°C from here onwards. ~50 μL of cells were dropped from a height of 30 centimeters above an ice cold, dry glass slide tilted at an angle of 45°C. Slides were air dried overnight and stored at 4°C. For chromosomes analysis slides were rinsed in PBS, dehydrated in 70%, 95%, 100% ethanol for 5 minutes and allowed to dry completely. Slides were denatured at 80°C on heat block with a 70% formamide, 2xSSC solution for 2 min and dehydrated with 70%, 95%, 100% ethanol for 5 min each. Once completely dry, slides were mounted with Vectashield antifade mounting medium containing DAPI (Vectashield, H-1200) and analyzed with EVOS FL AUTO Imaging System (ThermoFisher, AMAFD1000). Thirty independent metaphase spreads were analyzed for the parental mESCs and *hHPRT1* clone 1 and 25 independent spreads for *hHPRT1* clone 2.

## Acknowledgements

We thank Michael Kyba for helpful discussion and Haiping Hao and his team at the Johns Hopkins Deep Sequencing & Microarray Core Facility for Pacific Biosystems sequencing. This work was supported by NHGRI CEGS award RM1HG009491 and DARPA contract HR0011-16-2-0002 to J.D.B. T.D. and N.B. were supported by NCI award K99CA212621. J.D.B is a founder and director of the following: Neochromosome, Inc., The Center for Excellence for Engineering Biology, and CDI Labs Inc., and serves on the Scientific Advisory Board of the following: Modern Meadow, Inc., Recombinetics, Inc., and Sample6, Inc. L.A.M. is a founder of Neochromosome, Inc. All arrangements are reviewed and managed by the Committee on Conflict of Interest at NYU Langone Health. J.D.B. and L.A.M. are listed as inventors on patent application WO2018009809A1 describing eSwAP-In.

