## Supplemental data for "*De novo* assembly, delivery and expression of a 101 kb human gene in mouse cells"

**Supplemental information**

A.

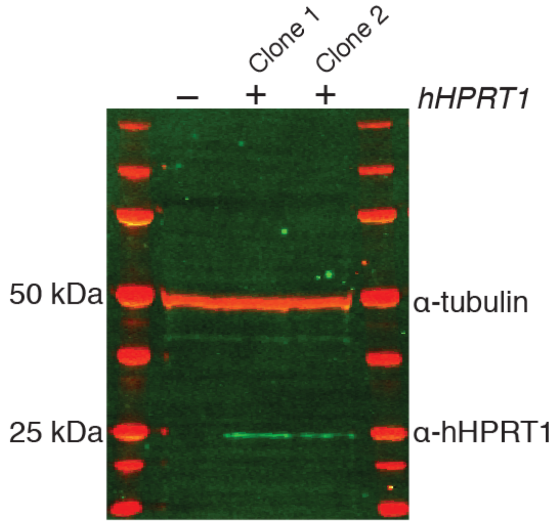


B.

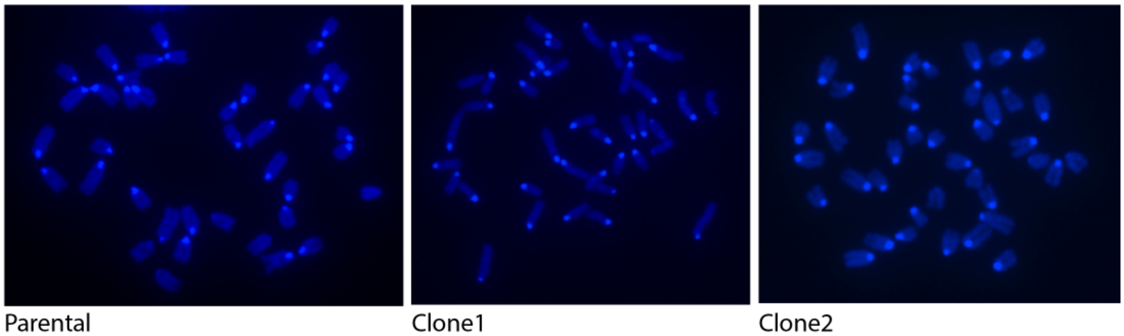


**Supplemental Fig. 1. *hHPRT1* clone characterization. (A)** Western blot analysis for *hHPRT1* protein expression in mouse ES cells using a human HPRT-specific monoclonal antibody. Parental mES (–) and *hHPRT1* clones 1 and 2 (+) were used. **(B)** Representative metaphase spreads from parental mESCs and *hHPRT1* clones 1 and 2.

### Supplemental Table 1: List of primers used to produce *HPRT1* amplicons

| For primer | Sequence | Rev primer | Sequence |
|---|---|---|---|
| HPRT.F 1 | gcggccgcTCCCATTACCATGGACAGTGTGCTT | HPRT.R 1 | gcggccgcGACCAGCATAATTCAGTACCCATGTT |
| HPRT.F 2 | gcggccgcGGTCAGTTACATGAAACAAGATGAAGTCA | HPRT.R 2 | gcggccgcTGTTTGTAATAGTTTCTAGGTACTATAATGGGA |
| HPRT.F 3 | gcggccgcGTCCATTAATTACTTCTTGATACTACTGACCT | HPRT.R 3 | gcggccgcGGGCTTGGAATCTTGACTAAAATTCTCC |
| HPRT.F 4 | gcggccgcCAGTAATATCCAGGGGTAATGGAGG | HPRT.R 4 | gcggccgcCAGTCACAAGTTAAGGGTGAAAACA |
| HPRT.F 5 | gcggccgcGAAGGAAAGAGTACAAACTATTTGGTTCAG | HPRT.R 5 | gcggccgcGCACAATTCCTTCCTTGGATGGT |
| HPRT.F 6 | gcggccgcATGATTCTGGGATACAAAATCCCTGG | HPRT.R 6 | gcggccgcTACAGTATCTTACACCCAAACCATCT |
| HPRT.F 7 | gcggccgcCCTAATTATTTCTAATACATGTCCTACCCC | HPRT.R 7 | gcggccgcGCTTAAATAATGCACCCTATCATTATTTAG |
| HPRT.F 8 | gcggccgcTCTTACATTGAGAAAAAACTGTATTTTCAAT | HPRT.R 8 | gcggccgcCTTGTGTGTGTACTTTTCAATTTAAGAAGTGA |
| HPRT.F 9 | gcggccgcCAGCTTCAGAATTCAACTTGCTTATAATTC | HPRT.R 9 | gcggccgcTACAGTGGTATAAGGTATGGGGACG |
| HPRT.F 10 | gcggccgcGTATGGCCTTAGACTATTCAGAACGTTA | HPRT.R 10 | gcggccgcatggggtgtaaaattgaaccataattgg |
| HPRT.F 11 | gcggccgcgggtcacagggcaagactttg | HPRT.R 11 | gcggccgcACAGTACAGTCAGCAAATGGGA |
| HPRT.F 12 | gcggccgcGAAATGGTGTTGCTGGAGCAAC | HPRT.R 12 | gcggccgcTTGGCAGAATTAAGTAAGTTGATGTTT |
| HPRT.F 13 | gcggccgcctcGAGTAATACCAGTTAAAAAATAGGCTAC | HPRT.R 13 | gcggccgcGAATGGGCAGAAATTGCTAGTTGG |
| HPRT.F 14 | gcggccgcCACAAATTATCCAAGGAGATGGCG | HPRT.R 14 | gcggccgcATCTACCTTGATTAAGGCAGTGTGC |
| HPRT.F 15 | gcggccgcTATGCCTTTCCCACTAGATTTTAAGC | HPRT.R 15 | gcggccgcTATGCCAGATCCTCTAGGATTAAATGC |
| HPRT.F 16 | gcggccgcAGCAAGGTTTGTATTTTCCTAGAACTTG | HPRT.R 16 | gcggccgcCAATCCCAAATCCAAACAGCCT |
| HPRT.F 17 | gcggccgcGATGACACTTCATGAGTTGACTATAATAATC | HPRT.R 17 | gcggccgcGCATATAGTATACATGCATAGCCAGTG |
| HPRT.F 18 | gcggccgcGTATTGAATGCTTGCATTGTATGTCT | HPRT.R 18 | gcggccgcGCTTTGCTTATGTTTAAGATGTCATGC |
| HPRT.F 19 | gcggccgcATAAGATCAATTCTGAGTGGTAGAAATGc | HPRT.R 19 | gcggccgcCACCATAATGCAGACTAATTTTCCCTC |
| HPRT.F 20 | gcggccgcCCAAAAAAACAACAAAGCCTCAGGA | HPRT.R 20 | gcggccgcTCAACCATTCTCTGTCCTTTTAACTG |
| HPRT.F 21 | gcggccgcACCTGAGCATATGTCCTTTCATACTTA | HPRT.R 21 | gcggccgcCAAGGATATGTCCCTCAAAAGTCTAGC |
| HPRT.F 22 | gcggccgcCCATTGAATCTCCTGTAAGGGTTTTATTG | HPRT.R 22 | gcggccgcACTGCAGCCTATTGGTAGCCTA |
| HPRT.F 23 | gcggccgcTACTGGGACCTCATACAAATGGGA | HPRT.R 23 | gcggccgcGGTCCAAGTGAATTGAAGAGGAAA |
| HPRT.F 24 | gcggccgcatCACTGTCTAACAGCCTCTCTTt | HPRT.R 24 | gcggccgcAACAGCATAGGTAAGGTGAGGAG |
| HPRT.F 25 | gcggccgcGATTCCAAAGCGGGTGAGGAAG | HPRT.R 25 | gcggccgcAACCCAAATGCTGCCTGTTGA |
| HPRT.F 26 | gcggccgcGCCCTTTCCAGCTCTTTAAACATATA | HPRT.R 26 | gcggccgcCCAGTTTCACTAATGACACAAACATGC |
| HPRT.F 27 | gcggccgcAAGATACACTCCCCAAAAGTTACTGA | HPRT.R 27 | gcggccgcAGCCAGCAGAAAAATCTGAAGAG |
| HPRT.F 28 | gcggccgcAGTCCAGATGACTTGTACATTAAACAC | HPRT.R 28 | gcggccgcGTGCCGAATTTGGTTACTCCTTT |
| HPRT.F 29 | gcggccgcACCTGGCCTTTGGAACTTGG | HPRT.R 29 | gcggccgcATCAGGGGGAAATGTTATTTATCATGAA |
| HPRT.F 30 | gcggccgcaGTTTCTTCCAGGGTGGCTTCT | HPRT.R 30 | gcggccgcTGTCCTTTGCCTGTGTTTTTAGGAA |
| HPRT.F 31 | gcggccgcAGCACCACAAAGTTAGAGGTCAA | HPRT.R 31 | gcggccgcAAGCATTATATCAGAAACAGAAAAATAATCATC |
| HPRT.F | gcggccgcCTCACTGAATGAGGCAGGTAGC | HPRT.R | gcggccgcaagACTTATTTCTAGTATTTTCTTCAT |

| | | | |
|---|---|---|---|
| 32 | | 32 | GATCG |
| HPRT.F 33 | gcggccgcGGAGAGAGAAGGCATAAACAATATTAAAA | HPRT.R 33 | gcggccgcaaaGCTGAGGAGAAAAATAAAAAGAATAC |
| HPRT.F 34 | gcggccgcTTTGGAGAAACAGCCCACCACC | HPRT.R 34 | gcggccgcaatGGTATGGGAGAATTGGGTTA |
| HPRT.F 35 | gcggccgctgCTGTGGATCGTTCAGCATGT | HPRT.R 35 | gcggccgccACGAAGCATAAATTTCTCCACAAA |
| HPRT.F 36 | gcggccgcGGTTATGTTCACTTCATTTGGTTACAG | HPRT.R 36 | gcggccgcAAAGATAATAAAATACCTCACATCATGAAtg |
| HPRT.F 37 | gcggccgcacGGTGTCTTAAATCTTTATGTGTTTG | HPRT.R 37 | gcggccgcCCATAATTAAATGTCTGTGTTCTGGTC |
| HPRT.F 38 | gcggccgcgcttcactctaaaatgatTGGACCATg | HPRT.R 38 | gcggccgcCATGGTGAGCGAGGTGAGGC |

**Supplemental Table 2: Yeast strains used in this study**

| Yeast strain | Genotype | Description | Reference |
|---|---|---|---|
| BY4741 | *MAT***a** *LYS2 met15Δ0 ura3Δ0 his3Δ1 leu2Δ0* | Wild-type lab strain for yeast assemblies | (1) |
| yLM1227 | BY4741 + *HPRT1 step 1::URA3* | *HPRT1* assembly step 1, isolate 1 | This study |
| yLM1228 | BY4741 + *HPRT1 step 1::URA3* | *HPRT1* assembly step 1, isolate 2 | This study |
| yLM1229 | yLM1227 + *HPRT1 step 2::LEU2* | *HPRT1* assembly step 2, isolate 1 | This study |
| yLM1231 | yLM1228 + *HPRT1 step 2::LEU2* | *HPRT1* assembly step 2, isolate 2 | This study |
| yLM1234 | yLM1229 + *HPRT1 step3::URA3* | *HPRT1* assembly step 3, isolate 1 | This study |
| yLM1235 | yLM1231 + *HPRT1 step3::URA3* | *HPRT1* assembly step 3, isolate 2 | This study |

**Supplemental Table 3: Plasmids used in this study**

| Plasmid | Description | Length (kb) | Reference |
|---|---|---|---|
| pLM453 | eSwAP-In assembly vector | 11 | This study |
| pLM718, pLM719 | *HPRT1* step 1, recovered from yLM1227 and yLM1228, respectively | 46 | This study |
| pLM747, pLM749 | *HPRT1* step 2, recovered from yLM1229 and 1231, respectively | 82 | This study |
| pLM750, pLM751 | *HPRT1* step 3, recovered from yLM1234 and yLM1235, respectively | 112 | This study |
| pLM707 | ICE loxP-loxM cassette | 4.5 | (2, 3) |
| pLM854 | pLM453 + loxP-loxM cassette | 13 | This study |
| pLM848 | pLM718 + loxP-loxM cassette | 48 | This study |
| pLM881 | pLM747 + loxP-loxM cassette | 84 | This study |
| pLM886 | pLM750 + loxP-loxM cassette | 114 | This study |

**Supplemental Table 4: QRT-PCR primers**

| Target | Forward primer | Rev primer |
|---|---|---|
| Mouse *Gapdh* | AGAACATCATCCCTGCATCC | CACATTGGGGGTAGGAACAC |
| Mouse *Nanog* | TTTCAGAAATCCCTTCCCTCG | TGATGAGGCGTTCCCAGAAT |
| Mouse *Esrrb* | CAGGCAAGGATGACAGACG | GAGACAGCACGAAGGACTGC |
| Mouse *Tcl1* | AAATTCCAGGTGATCTTGCG | TGTCCTTGGGGTACAGTTGC |
| Mouse *Gata6* | TTGCTCCGGTAACAGCAGTG | GTGGTCGCTTGTGTAGAAGGA |
| Mouse *T* | GCTTCAAGGAGCTAACTAACGAG | CCAGCAAGAAAGAGTACATGGC |
| Mouse *Pax3* | GCAGCGCAGGAGCAGAACCA | GCACTCGGGCCTCGGTAAGC |
| Mouse *Hand1* | CCCCTCTTCCGTCCTCTTAC | CTGCGAGTGGTCACACTGAT |
| Human *HPRT1* | GACCAGTCAACAGGGGACAT | CCTGACCAAGGAAAGCAAAG |
